# Molecular Signatures of Attention Networks

**DOI:** 10.1101/2023.06.29.547064

**Authors:** Hanna Schindler, Philippe Jawinski, Aurina Arnatkevičiūtė, Sebastian Markett

## Abstract

Attention network theory proposes three distinct types of attention - alerting, orienting, and control - that are supported by separate brain networks and modulated by different neurotransmitters, i.e., noradrenaline, acetylcholine, and dopamine. Here, we explore the extent of cortical, genetic, and molecular dissociation of these three attention systems using multimodal neuroimaging. We evaluated the spatial overlap between fMRI activation maps from the attention network test (ANT) and cortex-wide gene expression data from the Allen Human Brain Atlas. The goal was to identify genes associated with each of the attention networks in order to determine whether specific groups of genes were co-expressed with the corresponding attention networks. Furthermore, we analysed publicly available PET-maps of neurotransmitter receptors and transporters to investigate their spatial overlap with the attention networks.

Our analyses revealed a substantial number of genes (3871 for alerting, 6905 for orienting, 2556 for control) whose cortex-wide expression co-varied with the activation maps, prioritizing several molecular functions such as the regulation of protein biosynthesis, phosphorylation, and receptor binding. Contrary to the hypothesized associations, the ANT activation maps neither aligned with the distribution of noradrenaline, acetylcholine, and dopamine receptor and transporter molecules, nor with transcriptomic profiles that would suggest clearly separable networks. Independence of the attention networks appeared additionally constrained by a high level of spatial dependency between the network maps. Future work may need to re-conceptualize the attention networks in terms of their segregation and re-evaluate the presumed independence at the neural and neurochemical level.

## 1. Introduction

Attention prioritizes the processing of goal-relevant over less relevant information which ensures adaptive behavior in information-rich environments (Cowan, 1999; Posner & Fan, 2008). Attention network theory distinguishes between three different types of attention: i) alerting attention, which is the initiation of a state of heightened alertness in anticipation of upcoming stimuli; ii) orienting attention, which is the shift of the attentional focus to prioritize information processing at a particular spatial location, and iii) control attention, which is the selective amplification of relevant aspects of a stimulus when irrelevant information is present (Posner & Dehaene, 1994; Posner & Petersen, 1990). At the brain level, each type of attention is supported by distinct and distributed set of cortical and subcortical regions: the three attention networks *alerting*, *orienting*, and *control* (Petersen & Posner, 2012).

A core tenet of attention network theory is the presumed independence of the three attention networks. This independence is thought to arise from dissociating effects of large neuromodulatory systems: alerting is modulated by noradrenaline, orienting by acetylcholine, and control by dopamine (Posner & Rothbart, 2007). While the relevance of all three neuromodulators in attention is well established (Noudoost & Moore, 2011; Robbins, 1997), the evidence for the assumed dissociation and specificity in regard to the three attention networks is less conclusive. Experimental work with the attention network test (Fan et al., 2002), a behavioral protocol to assess the efficiency of all three attention networks simultaneously, was unable to detect influences of the potent acetylcholine agonist nicotine on any of the three attention networks <u>(see McCormick, 2022, for review)</u>. Furthermore, direct manipulations of the noradrenergic system by blocking the noradrenaline reuptake impact orienting attention but not alerting and control (Reynaud et al., 2019), while indirect measure of locus coeruleus activity suggest an involvement of noradrenaline in all three attention networks (Gabay et al., 2011; Geva et al., 2013). Only for dopamine, the evidence is more favorable, with clear demonstrations of increased dopamine release during task performance (Badgaiyan & Wack, 2011) and dissociating effects in patients with Parkinson’s disease whose cardinal symptom is dopaminergic impairment (Yang et al., 2022). It needs to be noted, however, that many studies have used rather indirect approaches to the neuromodulators and that neuroimaging work has been able to detect neuromodulation at the brain level in the absence of behavioral effects (Ikeda et al., 2017; Thienel et al., 2009).

The distal causation of these neuromodulatory effects is likely of genetic origin (Green et al., 2008). In line with the behavioral genetics literature (Polderman et al., 2015), it has been shown that the behavioral efficiency of the attention networks is heritable (Fan et al., 2001). Several studies have attempted to implicate candidate polymorphisms on genes with direct relevance for the neuromodulatory systems in behavioral and neural markers of attention and have found some evidence for the hypothesized dissociation (Fan et al., 2003; Fossella et al., 2002; Rueda et al., 2005). The available evidence, however, is not conclusive (Green et al., 2008; Posner et al., 2014) and genetic association studies on candidate genes have been disputed more recently, despite their apparent face value (Border et al., 2019; Montag et al., 2020; S. R. Moore, 2017).

Given the paucity of supporting evidence for the molecular and genetic dissociation of the three attention systems in humans, we decided to revisit the hypothesis by utilizing two novel and complementary approaches. Based on attention network theory’s assumption that the three attention networks can be activated by the attention network test (Fan et al., 2002; Fan et al., 2005), we hypothesize that the cortex-wide activation profile of each attention network corresponds with the relative availability of relevant receptor and transporter molecules (approach 1) and the relative expression of relevant genes in brain tissue (approach 2). Assuming that the molecular neuromodulation of attention networks occurs at the synaptic levels, we hypothesize that (1) brain regions that activate during alerting show higher availability of the noradrenaline transporter and higher expression of genes involved in noradrenaline binding, (2) that brain regions that activate during orienting show higher availability of nicotinergic acetylcholine receptors and vesilcular acetylcholine transporters, and a higher expression of genes involved in acetylcholine-gated cation-selective channel activity, and (3) that brain regions that activate during attention control show higher availability of the dopamine transporter and dopamine D1 and D2 receptors, and higher expression of genes involved in dopamine binding. We expect stronger covariation between task-evoked activity within each attentional domain and the mentioned gene sets as compared to randomly defined gene sets of equal size. We also expect stronger covariation between task-evoked activity within each attentional domain and receptor / transporter availability as compared to random null models of the attention networks. Moreover, we expect specificity of the associations in a way that gene sets and receptor / transporter availability with presumed relevance for one attentional domain is more strongly related to this domain than to the other two domains. Finally, given the assumed independence of the three attention networks, we expect (4) that the genome-wide transcriptomic signatures of the attention networks are uncorrelated.

To this end, we utilize a sample of healthy volunteers who completed the attention network test during functional magnetic resonance imaging (Markett et al., 2022), publicly available maps of group-level receptor and transporter distribution maps from positron emission tomography (Hansen, Shafiei, et al., 2021), and microarray measures for over 20,000 genes measured at 3702 locations in the brains of six donors as provided by the Allan Human Brain Atlas (AHBA) and curated by the abagen toolbox (Hawrylycz et al., 2012; Markello et al., 2021).

## 2. Methods

### 2.1 Functional imaging data set

We used the publicly available (https://osf.io/st9ae/) group-level attention network maps from our previous study (Markett et al., 2022). The maps were obtained from N = 78 healthy young volunteers (n = 35 female, n = 43 male, mean age M = 26.18, SD=5.34) who completed the revised attention network test (see next paragraph for details) during fMRI. Image-acquisition and preprocessing relied on pulse sequences and preprocessing routines from the Human Connectome Project (Glasser et al., 2013; Harms et al., 2018). Individual and group-level analyses were run on cifti-surface data after registration through multimodal surface matching (MSMsulc, Robinson et al., 2018) and minimal surface-smoothing (Gaussian filter with 4 mm width at half maximum). For details on image processing, we refer to our previous publication (Markett et al., 2022). All participants provided informed written consent and received remuneration. The study protocol adhered to the Declaration of Helskinki and was approved by a local ethics committee.

### 2.2 Attentional Network Test

The maps were derived by using the ANT (Xuan et al., 2016). The task followed a 4 x 2 design with the factors cueing condition (no cue, double cue, valid spatial cue, invalid spatial cue) and target (congruent flanker, incongruent flanker). Participants responded to a total of 288 trials, split into four sessions of 78 trials each. A typical trial sequence is shown in figure 1. Throughout each trial, participants were instructed to maintain fixation on a central fixation cross. Target stimuli appeared in one out of two boxes presented laterally to the fixation cross and included five arrows pointing either to the left or to the right. Participants were instructed to indicate as fast and as accurate as possible via button press whether the most central of the five arrows pointed left or right while ignoring the four flanking arrows that could either point into the same (congruent) or opposite direction (incongruent). Shortly before target onset, one out of four different cues was presented that carried information that a target was about to appear (temporal double cue) or where the target was about to appear (spatial cue). Spatial cues were either valid (i.e. giving correct information on the target location) or invalid (i.e. pointing at a location where the target did not appear). Some targets were preceded by no cue as a baseline condition. Cues were presented for 100 ms, the onset asynchrony between cues and targets was 0, 400, or 800 ms, targets were presented for 500 ms with an additional response window of 1200 ms, and trials were spaced with a jittered interval of 4,000 ms on average, systematically sampled from a distribution ranging from 2,000 to 12,000 ms. There were 48 trials with double cue, 48 trials with invalid cue, 48 trials with no cue, and 144 trials with valid cues. We realized an equal proportion of congruent and incongruent targets (144 each). The combinations of cue-target asynchronies, target location, and flanker types was counterbalanced for each cue condition. For more details on stimulus dimension and timing we refer to our previous work (Markett et al., 2022).

**Figure 1.**
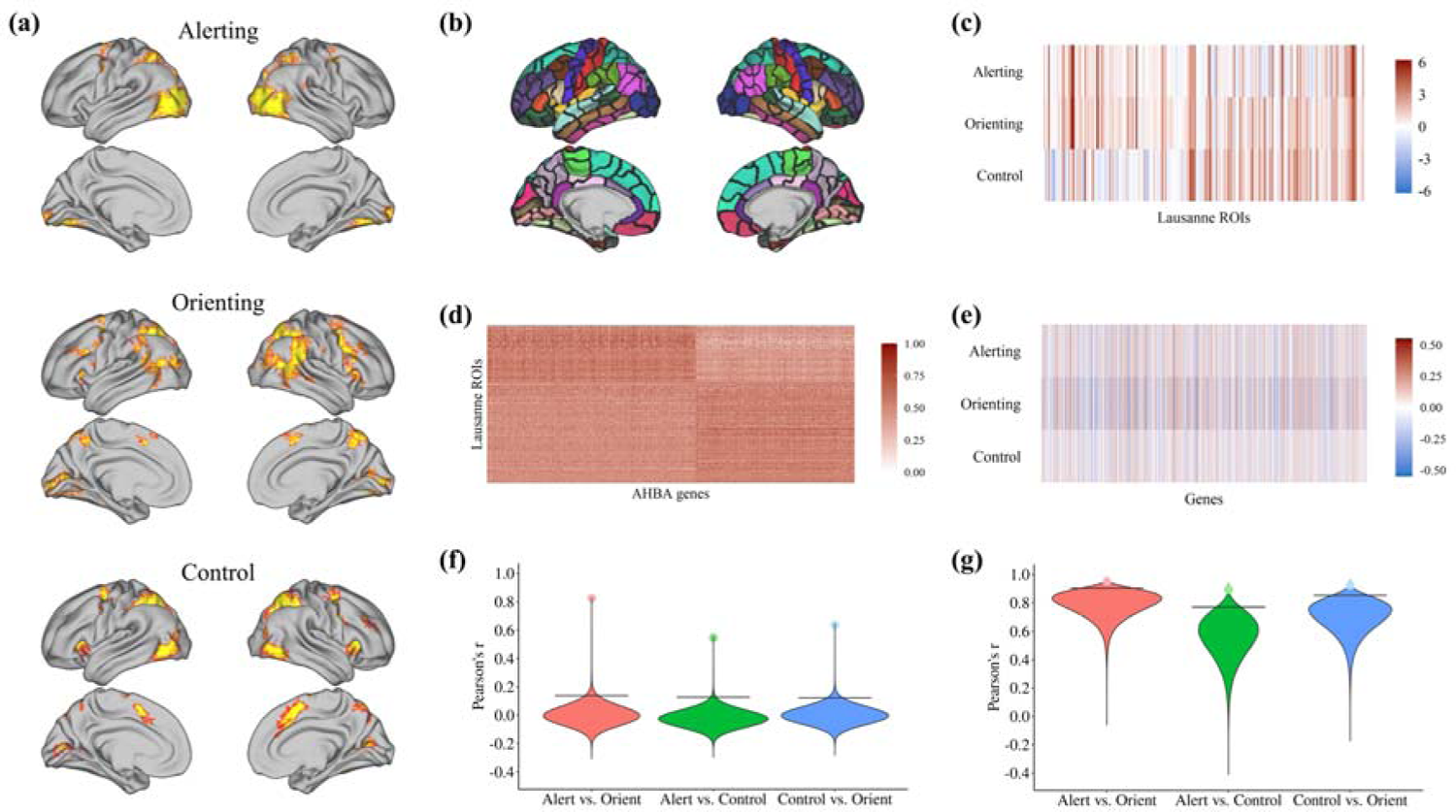
Illustration of the workflow how attention network images. (a) were decomposed into activation vectors (c) using Lausanne parcellation (b). Activation vectors (c) and abagen gene expression matrix (d) were then correlated with each other; correlation coefficients are represented in panel(e). (f) Violine plots of correlations between brain activation maps show the observed correlations (dots) of ANT maps vs. the distribution of expected correlations (violines) after one-million bidirectional random rotations of ANT maps. Horizonal lines reflect the 95th quantiles of correlations expected under the null. (g) Violine plots of correlations between gene associations results showing the observed correlations (dots) of gene association results vs. the distribution of expected correlations (violines) after one-million random rotations of ANT maps. Again, horizonal lines reflect the 95th quantiles of correlations expected under the null.

### 2.3 Attention Network Maps

The three attention networks were operationalized through linearly weighted contrasts on estimated beta images from first-level analyses in SPM12: the alerting network was obtained by contrasting the double cue minus the no cue condition across target conditions. The orienting network was defined as the validity effect by contrasting the invalid cue minus valid cue condition across targets. The control network was obtained by contrast incongruent minus congruent targets across cue conditions. Group-level maps were obtained by submitting the individual contrast images, using the Sandwich Estimator (SwE) Toolbox for SPM12 <u>6/29/2023 8:07:00</u> AM. Specifically, we implemented a modified SwE procedure that included a type c small sample size correction and a wild bootstrapping procedure consisting of 999 bootstraps.

### 2.4 Cortical parcellation

Linking imaging data from different modalities (functional activation maps, PET maps, gene-expression data) requires a brain parcellation as a common reference scheme. We utilized the Lausanne-219 parcellation which is a high resolution derivative of Freesurfer’s surface-based Desikan-Killiany-Atlas (Cammoun et al., 2012; Desikan et al., 2006) with 219 cortical brain regions. PET-maps were already available in this format. For the functional and gene-expression analyses, we created our own group version of the Lausanne-219 parcellation based on the T1-weighted structural images acquired alongside the functional imaging data. Structural images were run through HCP’s freesurfer pipeline (Glasser et al., 2013) and subsequently parcellated according to the Lausanne-219 atlas with scripts distributed with the CATO toolbox (de Lange & van den Heuvel, 2021). All individual parcellations were subsequently aligned with fsaverage_32LR space and a group atlas was created by assigning each vertex to the most frequent Lausanne-label across participants. We used this group atlas to extract region-wise mean activation differences from the unthresholded 2^nd^-level contrast maps for each ANT contrast. For the gene-expression analysis, we resampled the group parcellation to fsaverage5 space with 10k resolution for each hemisphere separately, as required by abagen.

### 2.5 Null model

Testing the spatial correspondence of different brain maps requires a null model that accounts for spatial non-independence (Alexander-Bloch et al., 2018). We evaluated statistical significance of the correlations between functional task activation and gene-expression profiles and between task activations and receptor/transporter maps through a spatial permutation approach (the “spin test”) where we swapped the parcellation labels randomly while accounting for the intrinsic geometry of the cortex (Váša et al., 2018). We created the null model based on centroid coordinates for each cortical region after projecting the group-level parcellation onto a sphere. For each permutation, coordinates were rotated around three axes with randomly generated angles. The same angles (with opposing signs) were used for both hemispheres to preserve hemispheric symmetry. To keep the alignment between rotated and unrotated parcellations intact, we matched each rotated to the closest unrotated region (minimum Euclidian distance), starting with the region that was most distant to all other regions on average and then progressing though all other regions in descending order. Code snippets for null model creations were taken from Github (github.com/frantisekvasa/rotate_parcellation).

### 2.6 Overlap of activation maps

We quantified the degree of spatial overlap between the phenotypic activation maps (brain images) by calculating product-moment correlations between respective activation vectors. Statistical significance was determined via permutation testing based on the null model. We randomly rotated each activation map one million times to obtain permutated activation maps that account for spatial autocorrelations in the brain maps (see section 2.5). For each pairwise comparison (alerting vs. control, alerting vs. orienting, and control vs. orienting), rotation was performed twice, i.e., one activation vector (e.g., alerting) was rotated first and correlated with the observed activation vector (e.g., control) and vice versa. Thus, each observed correlation (e.g., alerting vs. control) was compared to two million expected correlations (alerting observed vs. control rotated, and alerting rotated vs. control observed).

### 2.7 Gene expression analyses

For gene expression analyses, we used *abagen* (Markello et al., 2021), an open-source Python interface that enables integration of the Allen Human Brain Atlas (AHBA; Hawrylycz et al., 2012) with neuroimaging data. The AHBA is an open-access database containing gene expression and other microarray expression data collected from six human post-mortem brains. *abagen* intends to improve workflows related to the AHBA in terms of transparency, reproducibility, and standardization. The tool enhances transcriptional data analyses as these are often accompanied by a wide range of processing options making results from different studies less comparable.

We utilized *abagen*s workflow for correlated gene expression analyses, employing its default parameters. The detailed processing steps are documented in the *abagen* methods report (Appendix B). This procedure involved uploading a surface brain atlas to generate a genes-by-regions expression matrix with a total of 15632 incorporated genes.

### 2.8 Gene expression and attention networks

We correlated each of the three regions-by-activation vectors from the ANT conditions with the regions-by-genes expression matrix obtained from *abagen,* resulting in one product-moment correlation coefficient for each ANT condition by gene combination. We obtained corresponding p-values through permutation tests with the rotated spatial null model (see section 2.5). We performed 1 million rotations to determine significance with adequate accuracy even after stringent Bonferroni-correction for multiple testing (0.05/15632 = 0.0000032). For each gene, we determined the relative frequency by which the absolute correlation coefficient after random permutation was equal to or larger than the absolute empirical (observed) correlation coefficients. To distinguish positive from negative correlations, the p-values were transformed into z-values under consideration of the empirical correlations’ signs. In a last step, all genes were rank-ordered based on these z-values. Rank ties were resolved by inspecting the empirical correlation coefficient, allowing duplicate z-values to be unambiguously assigned to a rank. These rank indices, each obtained by considering both the p-value and empirical correlation coefficient with its sign, serve as input for the subsequent gene set enrichment analysis. In this analysis step, no thresholds were applied, and all genes, along with their respective rank indices, were preserved without exclusions.

### 2.9 Gene-expression-similarities of attention networks

We calculated pair-wise correlations between the three vectors containing the association between each gene by ANT condition combination (see section 2.8) to quantify gene-expression-similarities between the three attention networks. Corresponding p-values were computed by randomly rotating the ANT activation maps relative to the gene expression maps (see section 2.6) and calculating the proportion of pair-wise correlation coefficients of genetic associations results that were at least as extreme as the original pair-wise correlation coefficients. ANT activation maps were rotated in equal directions to preserve their phenotypic correlations. Under the assumption of no association between ANT activation maps and gene expression maps, the pair-wise correlations of association results were expected to reflect, on average, the phenotypic correlations between ANT activation maps. In case of true associations between ANT activation maps and gene expression maps, genetic similarities (i.e., pair-wise correlations of genetic association results) were expected to exceed the phenotypic correlation.

### 2.10 Gene set definition

We grouped the gene expression decoding results for each ANT conditions by their underlying molecular functions to detect similarities between single genes on a higher level. A common tool for classifying such gene functions is the PANTHER Classification System (Mi et al., 2021). PANTHER allows functional sorting of proteins, based on various criteria such as signaling and metabolic pathways or external aspects of the Gene Ontology database. The Gene Ontology (Ashburner et al., 2000; Gene Ontology Consortium, 2021) provides biological information to PANTHER that can be divided into three domains: Molecular Functions, Biological Processes, and Cellular Components (Mi & Thomas, 2009). Given our hypotheses on local activity of dopamine, norepinephrine, and acetylcholine, we focussed our analyses only molecular functions (MF), i.e., on all molecular activities that arise almost directly from one or more gene products, such as binding or transport activity. In contrast, biological processes represent activities superordinate to MF, thus not describing actions on a local, individual level but rather general metabolic or physiological processes. Cellular components, on the other hand, describe cell structures (e.g., mitochondria, cytosol) instead of cell activity and therefore do not provide relevant information about the neurotransmitter system.

Further, the selection of MFs we hypothesize being involved in alerting, orienting, and control focused primarily on transmitter interaction rather than synthesis since regions of transmitter synthesis are relatively confined. We rather expect the experimental manipulation leading to increased postsynaptic activity in the target regions of the three transmitter systems. Before performing statistical analyses, we pre-selected those three MF (out of around 5000 available MF in the database) that best reflect neurotransmitter binding activity of dopamine, norepinephrine, and acetylcholine:

1. Dopamine Binding (GO:0035240, 7 Genes) Binding to dopamine, a catecholamine neurotransmitter formed by aromatic-L-amino-acid decarboxylase from 3,4-dihydroxy-L-phenylalanine.
2. Norepinephrine Binding (GO:0051380, 4 Genes) Binding to norepinephrine, (3,4-dihydroxyphenyl-2-aminoethanol), a hormone secreted by the adrenal medulla and a neurotransmitter in the sympathetic peripheral nervous system and in some tracts of the CNS. It is also the biosynthetic precursor of epinephrine.
3. Acetylcholine-Gated Cation-Selective Channel Activity (GO:0022848, 18 Genes) Selectively enables the transmembrane transfer of a cation by a channel that opens upon binding acetylcholine.

### 2.11 Gene Set Enrichment Analysis (GSEA)

We used GSEA to assess whether genes belonging to the specified gene sets Dopamine Binding, Norepinephrine Binding, and Acetylcholine-Gated Cation-Selective Channel Activity showed stronger co-expression with the attention network maps relative to all other expressed genes. Association results derived from correlating gene expression maps and ANT activation maps (see section 2.8) were uploaded to PANTHER v17, including unique Gene IDs and their corresponding rank index. This index indicates the extent to which gene expression and fMRI activation spatially overlap in comparison to other genes. Identical tool settings and processing steps were applied to all gene lists (Appendix B). The results of these enrichment tests are lists of significantly over- or under-represented MFs for each ANT contrast as well as information about all mapped genes for each MF. GSEA was primarily applied to evaluate whether the three hypothesized categories are statistically overrepresented. In addition, we explored the potential over- or under-representation of other categories. However, it is crucial to interpret these exploratory findings with caution, considering the autocorrelations of gene expressions.

Interpretation of the enrichment results was facilitated by *GO-Figure!* (Reijnders & Waterhouse, 2021), an open-source Python software that clusters GO-terms and hence reduces redundancies. Functionally similar MFs are arranged in a cluster, followed by the selection of one representative MF. All representatives are then plotted in a two-dimensional space based on their pair-wise semantic similarity, which is a numerical value indicating the extent of functional, structural, and hierarchical similarity within the Gene Ontology. We included all significant MFs (p<.05, FDR corrected) when building summary visualizations with *GO-Figure*.

### 2.12 PET data set

We obtained 19 publicly available group-level maps of receptor and transporter availability in nine neurotransmitter systems (Hansen, Shafiei, et al., 2021), derived from PET imaging in 27 different samples with N = 1239 healthy participants in total. From the 19 maps, we selected six maps with direct relevance for the three hypothesized neurotransmitter systems: Noradrenaline (noradrenaline transporter, NET), acetylcholine (nicotinergic receptor a4b2 and vesicular acetylcholine transporter, vACht), and dopamine (d1-receptor, d2-receptor, and dopamine transporter, DAT). All data were already provided in the Lausanne-219 parcellation (github.com/netneurolab/hansen_receptors). Relationships between attention networks and PET maps were assessed through Pearson correlations. Corresponding p-values were obtained through permutation testing with the spin test (5000 permutations each). Given that the statistical tests on the six transporter and receptor maps were hypothesis-guided tests, we set the statistical threshold to p<.05, uncorrected. We also ran an exploratory analysis on correlations between ANT maps and the other 13 PET maps. In the absence of a hypothesis, p-values were adjusted with a false discovery rate of q=.05.

### 2.13 Open Science Statement

Code for structural and functional preprocessing can be obtained from https://github.com/Washington-University/HCPpipelisnes. Group-level data, decoding results, and analyses code can be obtained from github.com/schinhan/ant_genes. We have published other work on the same data set (Markett et al., 2022).

## 3. Results

### 3.1 Overlap of activation maps

In a first step, we assessed the spatial overlap between the three attention networks. The spatial activation vectors of attention networks were positively intercorrelated: r = .546 (p = 3.5e-06) for alerting vs. control, r = .824 (p = 2e-06) for alerting vs. orienting, and r = .636 (p = 1e-06) for control vs. orienting. P-values indicate that, for each pairwise comparison, almost none of the randomly rotated brain maps exhibit stronger correlations with the observed brain maps than the observed brain maps themselves (Fig. 1f).

### 3.2 Gene Expression Analyses

Secondly, we examined the spatial correspondence between gene expression and spatial activation of attention networks for each gene in comparison to all other genes. The gene expression matrix extracted from *abagen* is shown in Figure 1d. Correlation of the region-by-gene expression matrix with the spatial activation vectors of the ANT conditions (Fig. 1c) yielded an ANT condition-by-gene association matrix (Fig. 1e). Significance testing with one million permutations of the spatial null model revealed 3871 genes whose cortex-wide expression patterns covaried with the activation maps (p<.05, FDR corrected) in alerting, 6905 genes in orienting, and 2556 genes in control. Association results of all expressed genes and the different ANT conditions are displayed in respective Manhattan plots (Fig. 2a-c), with further details provided in Table B1 (Appendix B). We note that Manhattan plots do not highlight specific genomic regions that contain accumulations of strong association p-values as typically shown in genome-wide association studies. In this expression-based analysis, clusters of genes may span the whole genome and still overlap in their cortex-wide expression patterns, producing ‘horizontal band of associations’ with similar test-statistics in Manhattan plots (e.g., see band of associations in Fig. 2b ranging from -log_10_(*p*) = 4 to -log_10_(*p*) = 5). To identify groups of genes with a relative enrichment of signals, we carried out Gene Set Enrichment Analyses (GSEA), which implies that all genes (both significant and non-significant genes) were included along with their corresponding z-values.

**Figure 2.**
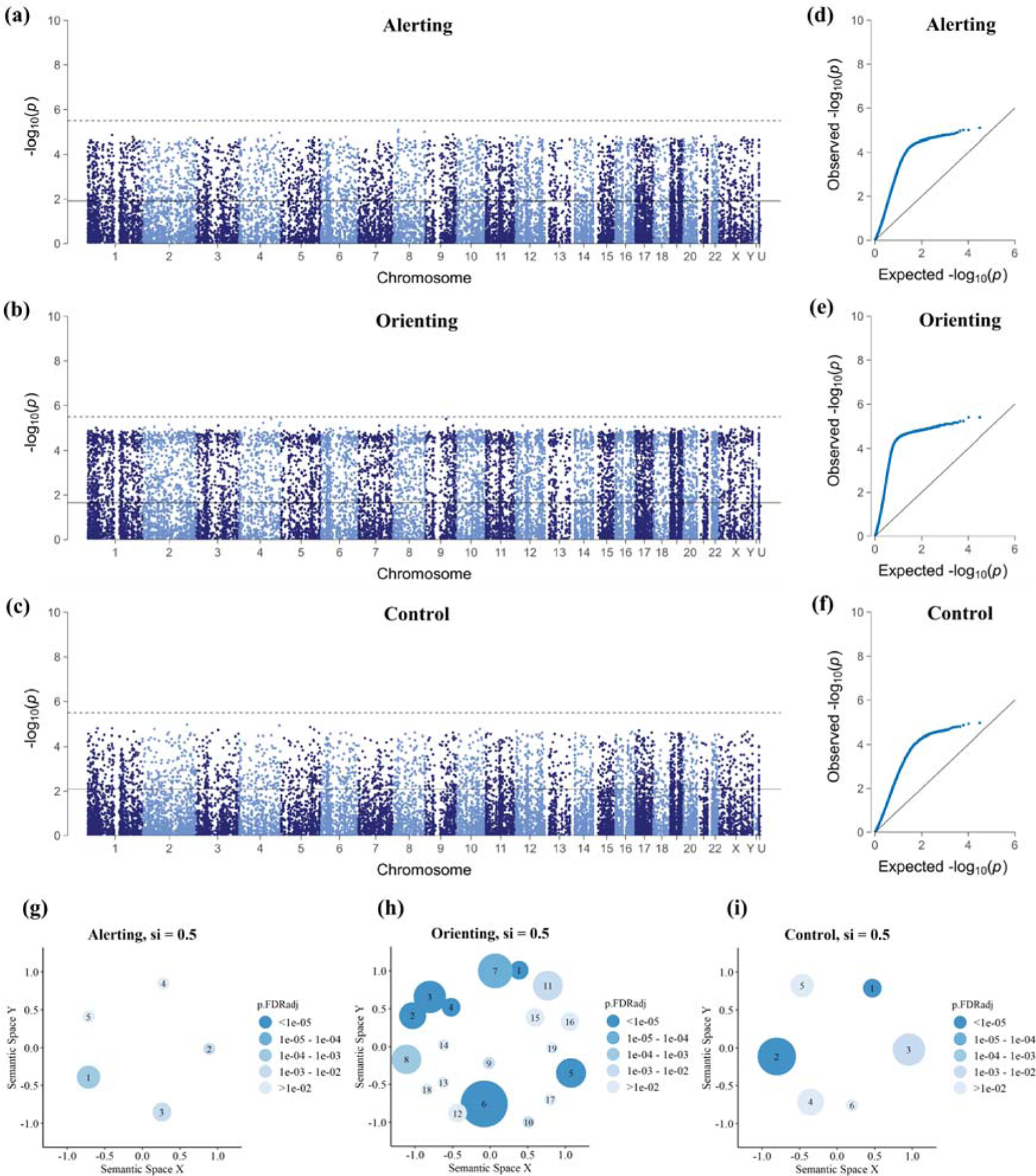
Summary of single-gene and GSEA results for the three ANT conditions. Manhattan plots (a–c) show the genomic position of genes on the x-axes (autosomes in ascending order, gonosomes X and Y, genes in category U remained unassigned in the matching of genes to chromosomes). The y-axes show p-values (-log10 scale) obtained through permutation tests (see *section 2.8*). The lower horizontal line marks the FDR-corrected level of significance, the upper dashed horizontal line marks the Bonferroni-adjusted level of significance. QQ-plots (d-f) compare the distribution of observed p-values (y-axes) against the distribution of expected p-values under the null hypothesis (y-axes). Leftward deflections (blue dots) from the projected null (diagonal line) indicate an enrichment of low p-values. GO-Figure! plots (g-i) summarize enriched MF from GSEA. Cluster colors represent the p-value of the selected representative. The circles size indicates the number of MF assigned to the cluster. Spatial distances between clusters represent their semantic similarity. Labels of the cluster’s representatives are displayed in *Table 2*.

**Table 2.**
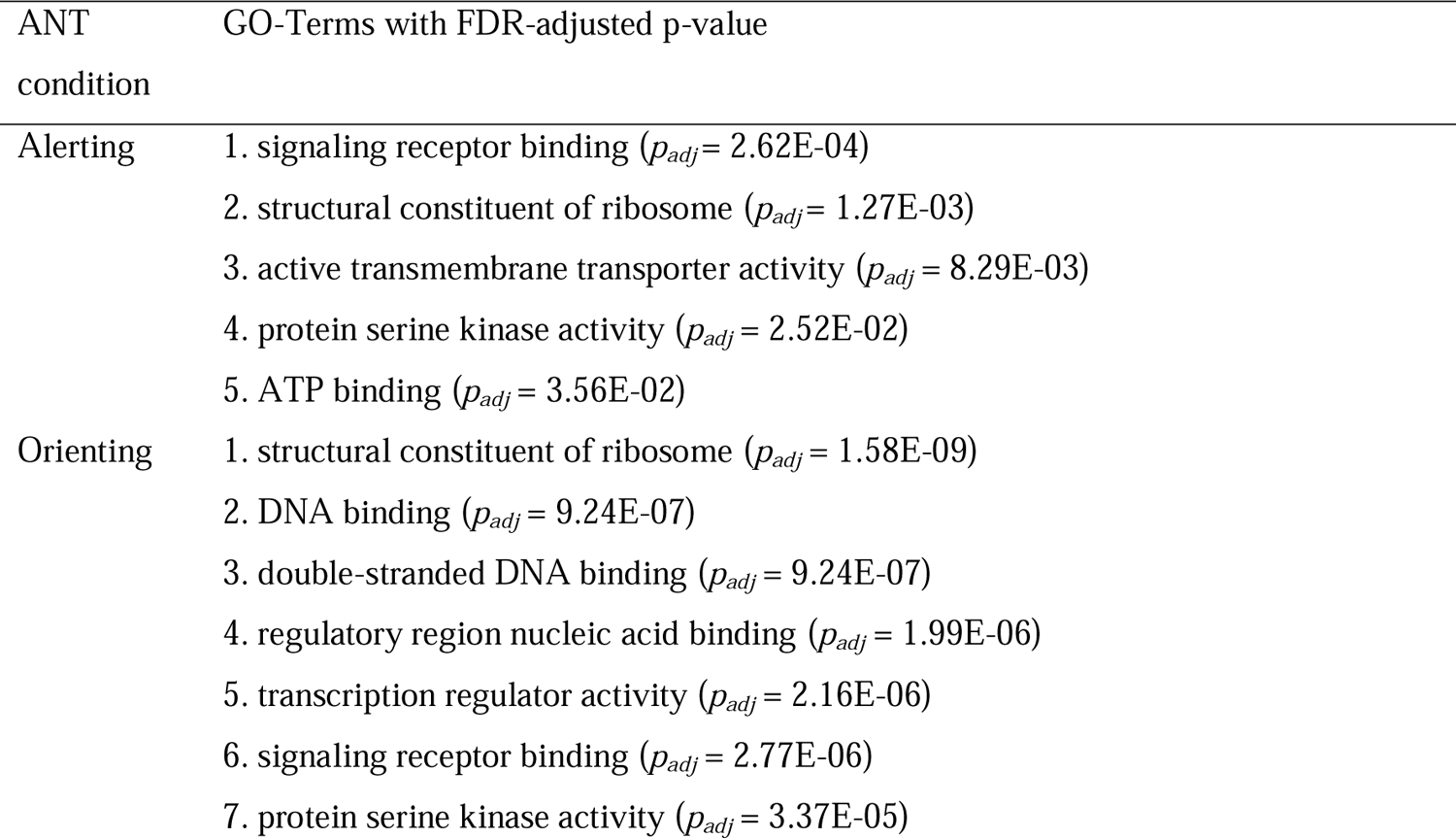
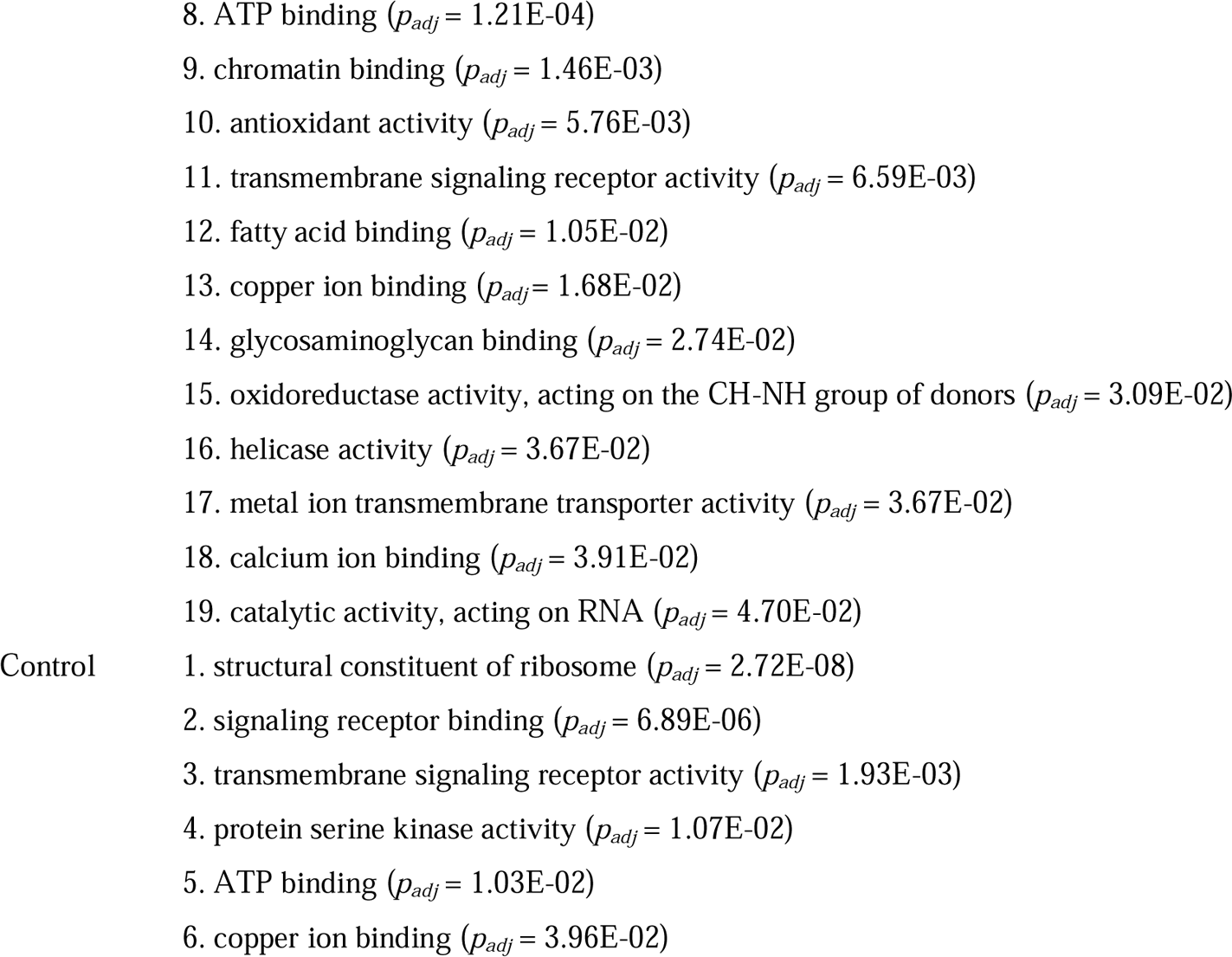
GO-Terms functioning as representatives of clusters generated with GO-Figure (si=0.50).

### 3.3 Gene-expression-similarities of attention networks

Next, we aimed to test whether the general gene expression patterns showed spatial overlap between the alerting, orienting, and control network.The empirical correlation between vectors containing the genetic associations of ANT maps, which, again, represent the correlation coefficients between activation maps and gene expression maps, aggregated to r = .893 (p = .0002) for alerting vs. control, r = .948 (p = .0015) for alerting vs. orienting, and r = .922 (p = .0008) for control vs. orienting. By comparison, under the null model of no association between gene expression and ANT activation maps (operationalized through one million random rotations of the ANT activation maps), correlations of the gene association results derived for the three ANT conditions were lower and approximated the direct correlations of ANT maps (mean r = .554 for alerting vs. control, mean r = .785 for alerting vs. orienting, and mean r = .695 for control vs. orienting). The enhanced similarity in association results as indicated by higher-than-expected correlation coefficients suggests a shared systematic covariation between gene expression and ANT maps (Fig. 1g).

### 3.4 Gene Set Enrichment Analysis

GSEA were applied for all three ANT conditions to quantify the probability to which the genes associated with their hypothesized MF are coincidentally or systematically ranked higher within the whole gene list. To do so, we uploaded association results of all 15632 genes to PANTHER, of which 14466 genes could be successfully annotated within the database. Consequently, we derived GSEA results from these 14466 genes and excluded the unmapped gene IDs from subsequent analysis.

### 3.4.1 A priori Hypotheses

Table 1 shows enrichment analysis results (Mann-Whitney U Test Statistic) for all three MF defined a priori. Test statistics inform about how many genes were mapped to this MF and whether the MF is over- or underrepresented. No ANT contrast was significantly positively related to its matching MF: There was no evidence for positive enrichment of genes related to norepinephrine binding in alerting, nor for dopamine binding in control, nor for acetylcholine-gated cation-selective channel activity in orienting. Furthermore, there was also no positive enrichment for any pre-defined MF and the other two ANT conditions. In contrast, we did find an enrichment towards the bottom extreme for norepinephrine binding and the control network, meaning that the distribution of values of this gene set (z scores derived from the permutation-based p-value and the sign of the observed correlation coefficient) was shifted towards smaller values relative to the overall list of genes, suggesting lower expression at activated regions.

**Table 1.**
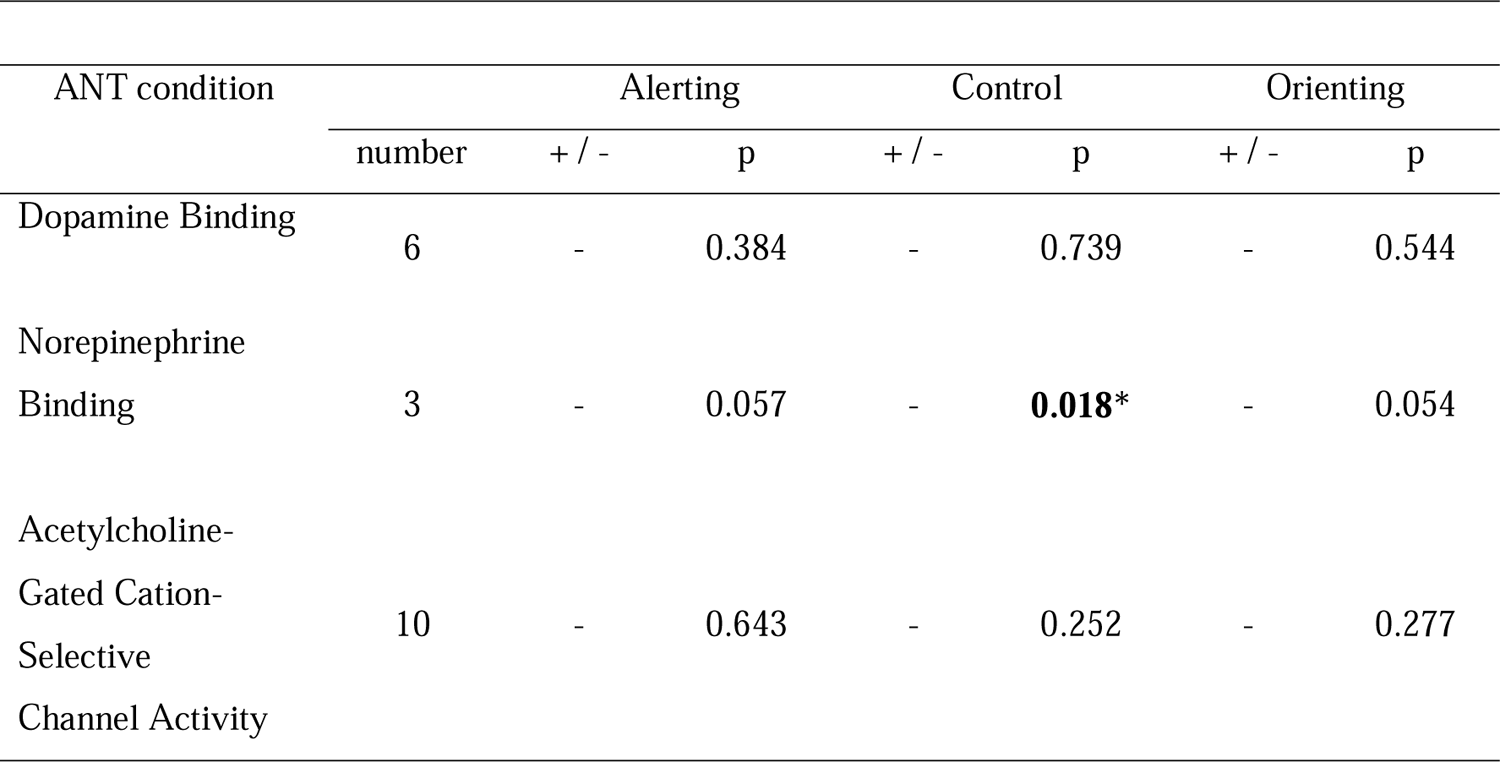
GSEA results of MF defined a-priori.

| ANT condition | Alerting |  |  | Control |  | Orienting |  |
| --- | --- | --- | --- | --- | --- | --- | --- |
|  | number | + / - | p | + / - | p | + / - | p |
| Dopamine Binding | 6 | - | 0.384 | - | 0.739 | - | 0.544 |
| Norepinephrine Binding | 3 | - | 0.057 | - | **0.018\*** | - | 0.054 |
| Acetylcholine-Gated Cation-Selective Channel Activity | 10 | - | 0.643 | - | 0.252 | - | 0.277 |

### 3.4.2 Exploratory Findings

Even though the pre-selected MF did not yield any significant hits, we found positive enrichment (p<.05, FDR-corrected) for 4 other MF in alerting (Table C.1, Appendix C), 41 MF in orienting (Table C.2), and 8 MF in control (Table C.3). Negative associations appeared for 4 MF in alerting, 25 MF in orienting, and 17 MF in control. Among these hits, no MF was directly related to dopamine, acetylcholine, or norepinephrine (see Appendix C).

Figure 2 (g-i) displays the summarized visualisation, implemented using *GO-Figure* (see Appendix D for detailed descriptions). The GO-terms of all significant MF (p<.05, FDR corrected) with the corresponding p-value served as input. As parameters, we set a maximum of 20 clusters, a default similarity cut-off of si=.50, and restricted the analysis to molecular functions. The similarity cut-off is a parameter that is set between 0 and 1 determining the degree of semantic similarity necessary for two GO-terms within the Gene Ontology to be assigned to the same cluster. The selected GO-Terms functioning as representatives of all clusters are listed in table 2.

### 3.5 Attention networks and neurotransmitter systems measured by PET

Our complementary methodological approach involved the analysis of PET-maps representing neurotransmitter receptors and transporters to evaluate their spatial overlap with the attention networks. Based on attention network theory, we expected brain regions with stronger activation during attention to correlate with the availability of receptor and transporter molecules within three neurotransmitter systems. As shown in figure 3, however, empirical results were not in line with our hypotheses: The availability of the noradrenalin transporter did not match the alerting network, and the availability of dopamine D1 and D2 receptors and the dopamine transporter did not correspond with the control network. Also, the availability of the most abundant nicotinergic acetylcholine receptor did not correlate with the activation of the orienting network. Only the availability of the vesicular acetylcholine transporter showed nominally significant correlations with the orienting network. The relationship, however, was negative, indicating that brain regions with higher transporter availability were less active during orienting of attention (see figure 3), which suggests that the acetylcholine transporter is not overexpressed within the orienting network. The exploratory analyses (13 tests, FDR-adjusted) on other neurotransmitter systems did not reveal significant relationships, except for a negative correlation between dopamine D1 availability and the alerting network (see figure 3).

**Figure 3.**
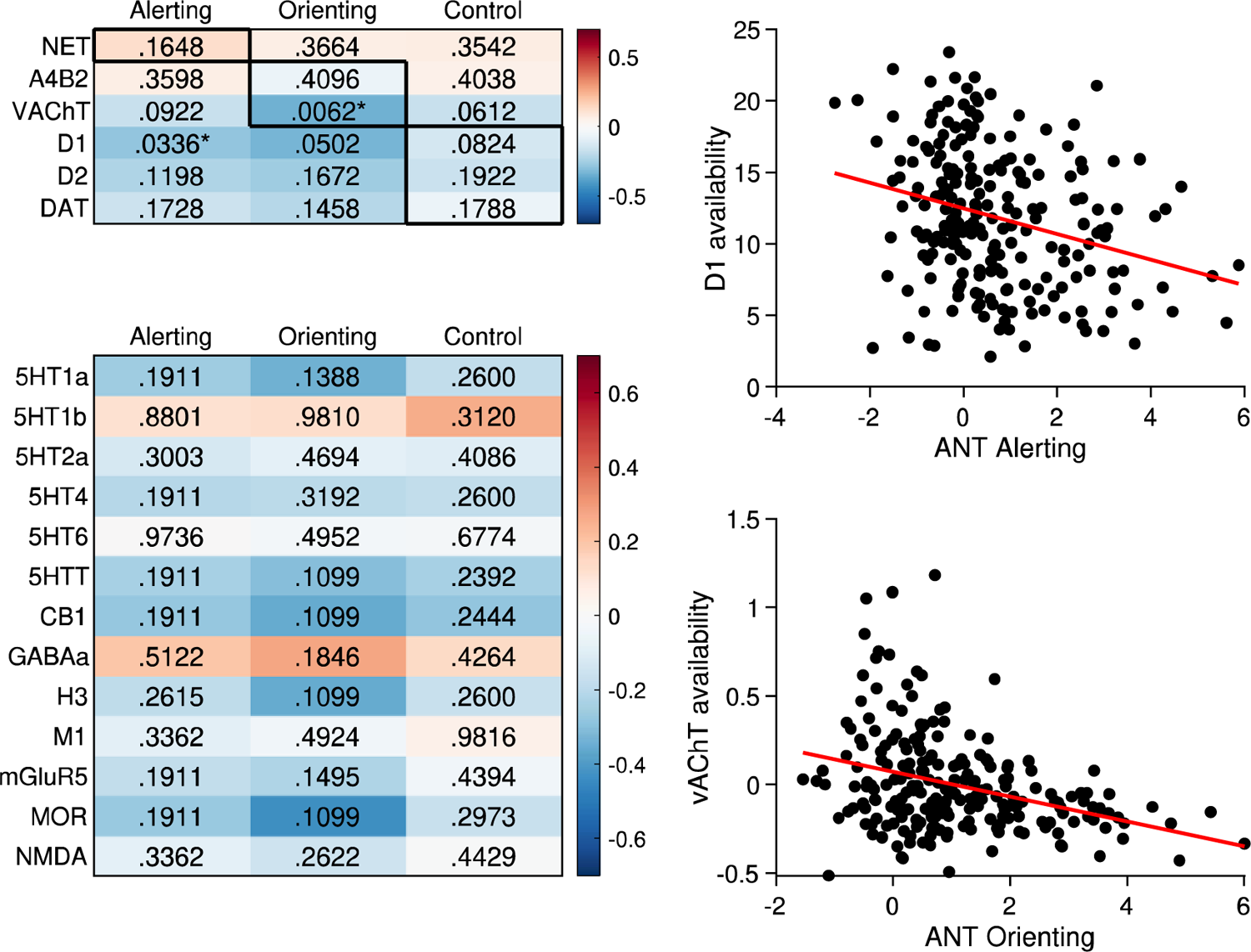
The left-hand panel shows correlations between neurotransmitter maps as revealed by PET-imaging (rows) and attention networks (columns) based on the Lausanne-219 parcellation. P-values were obtained through permutation testing under consideration of the autocorrelations across brain regions. The upper part of the table shows the hypothesized relationships (bold-framed cells), the lower part of the table gives the results from the exploratory analysis. Significant correlations are marked with an asterisk. The right-hand panels visualize the two significant correlations between regional dopamine D1 availability and activity in the alerting network (top) and regional availability of vAChT an neural activity in the orienting network (bottom).

## Discussion

Attention network theory states that three distinct attention networks are each modulated by a specific neurotransmitter: the alerting network by norepinephrine, the orienting network by acetylcholine, and the attention control network by dopamine. We sought to ascertain these hypotheses by assessing the molecular signatures of attention networks through a multimodal neuroimaging design: We compared fMRI activation maps from the attention network test with cortex-wide gene expression patterns and with the spatial distribution of neurotransmitter receptors and transporters as revealed by PET imaging.

If the three attention networks were modulated by distinct neurotransmitters as proposed by attention network theory, we would expect spatial correspondence between the networks’ activation patterns on the one hand and the expression-levels of genes related to transmitter binding as well as the availability of receptor and transporter molecules of the proposed neurotransmitter on the other hand.

### No evidence for the hypothesized neuromodulators

We did not find any evidence for the suggested neuromodulatory separation for any of the three attention networks. Even though a substantial number of genes co-expressed significantly with the attention networks, there was no enrichment of gene sets linked to the molecular functions of dopamine binding, norepinephrine binding, and acetylcholine-gated channel activity. Regarding the spatial distribution of receptor and transporter molecules in the PET images, there was also no spatial correspondence with attention network in the hypothesized direction: the distribution of NET did not correlate with the activation of the alerting network, the distribution of a4b2 receptors did not correlate with activation of the orienting network, and neither DRD1 nor DRD2 availability correlated with the activation of the attention control network. Only the distribution of the vesicular acetylcholine transporter correlated with the activation of the orienting network, however, in the opposite direction, suggesting higher transporter availability outside the orienting network.

From this, we conclude that the specific hypotheses on distinct neuromodulation of the three attention networks do not hold, at least not at the level of transcriptomic activity and the availability of receptor or transporter molecules, and at least not in the general and straightforward manner as suggested by attention network theory. We will discuss implications against methodological questions further below.

### No evidence for a transcriptomic separation of attention networks

Despite the unfavorable evidence for the a priori hypotheses, our exploratory analysis revealed that our design was at least in principle capable of detecting molecular functions in relationship with the attention networks. We found a substantial number of genes (3871 for alerting, 6905 for orienting, 2556 for control) whose cortex-wide transcription co-varied with the activation maps. Among a ranked list of all available genes, gene set enrichment analysis further prioritized several molecular functions for all three attention networks. These included genes involved in the regulation of protein biosynthesis, phosphorylation, enzymatic activity, and receptor binding. The implication of such broad terms in all three networks is not surprising, given the high similarity of the co-expression patterns across networks. Such broad terms, however, could also result from spatial autocorrelations of the gene expression maps(Fulcher et al., 2021), and should therefore be interpreted with caution.

In absolute numbers, we implicated most genes in the orienting network. The additional molecular functions prioritized here included genes involved in transcriptomic activity and regulation. Among all identified gene sets, the only gene set with direct relevance for a concrete neurotransmitter was glutamate receptor activity (GO:0008066). This gene set, however, was underrepresented among the associations for the control network. In sum, the exploratory analysis of the attention networks’ transcriptomic signatures revealed a relatively broad set of associated molecular functions. Contrary to the claims of attention network theory, there was no clear distinction in the networks’ transcriptomic profiles and no evidence for any given neurotransmitter system whose neuromodulatory activity might support the hypothesized separation of the networks at the molecular level.

### Methodological Considerations

The current approach is based on various assumptions of attention network theory. In the following, we will discuss the current finding against these assumptions and highlight some challenges within attention network theory that may prompt a need for reevaluation of its central principles.

One of the fundamental assumptions of attention network theory is the independence of the three attention networks. Independence refers to the idea that the three networks exhibit uncorrelated behavioral responses when manipulated, distinct activation patterns, and independent neuromodulatory influences (Petersen & Posner, 2012; Posner & Fan, 2008; Posner & Rothbart, 2007). While the behavioral independence of the three networks has been well-documented (Callejas et al., 2004; Fan et al., 2002, 2009; Ishigami & Klein, 2010; Markett et al., 2022), our findings suggest that there is significant overlap between the cortical activation patterns of the three networks. This overlap limits the scope of the theory but does not necessarily rule out the possibility of separate neuromodulation which may still occur despite overlapping activation patterns. This, however, was not the case. The lack of independence in functional activation, transcriptomic activity, and receptor expression, impose significant constraints on the predictions of attention network theory.

Our approach also rests on the assumption that the three attention networks can be operationalized using activation maps from the attention network test (ANT). This assumption is based on attention network theory, which views the ANT as the standard protocol for separating the networks and considers it capable of fully activating them (Fan et al., 2005; Fan & Posner, 2004). By keeping visual stimuli constant and by counterbalancing motor responses, the ANT is believed to reveal the extra neural effort required for different attention systems. But while the ANT’s validity can be reasonably assumed, it is unclear whether the presumed transmitter systems modulate a given network as a whole (and not only selected regions within the network) and if activation across the network is proportional to the hypothesized neuromodulation. Such proportional relationship would imply uniform neuromodulation of the several processes captured by the ANT activation maps such as increased activation due to the prioritization of information processing in sensory cortices (Brefczynski & DeYoe, 1999; Kastner et al., 1999; Müller et al., 2003), increased activation due to the maintenance and allocation of the attentional focus in premotor cortex and the frontal eye fields (T. Moore et al., 2003; Thompson, 2005), and increased activity linked to the detection of increased demand for attentional resources in the anterior cingulate (Botvinick et al., 2004). Since attention network theory does not specify the mechanisms of neuromodulation, the current results do not necessarily refute the specific hypotheses. However, they raise doubts that the attention networks as currently operationalized are modulated as a whole, and indicate the need to revise the theory. This revision should include a specification of the relationship between attention networks and activation patterns, the functional neuroanatomy of each network, and the potential target regions of neuromodulatory influences.

A further assumption of our approach is that the hypothesized neuromodulation can be operationalized using three gene sets from the gene ontology that encompass molecular functions involved in transmitter binding. Transmitter binding describes the process where neurotransmitters interact with synaptic molecules. The gene products of the tested genes included the dopamine transporter, dopamine receptors and associated intracellular signaling molecules, different adrenergic receptors and proteins forming subunits for the pentameric nicotinergic acetylcholine receptor. If the experimental manipulation within the attention network test would lead to the hypothesized activity increases of dopaminergic neurons in the midbrain, noradrenergic neurons in the locus coeruleus, and cholinergic neurons in the basal forebrain, we would expect increased neural activation in their projection areas. These projection areas are characterized by an abundance of the mentioned receptor and transporter molecules, which should also be apparent in their transcriptomic activity. Our present findings, however, suggest that there is no detectable relationship between neural activation patterns elicited by the experimental manipulation of attention and receptor or transporter availability, or related transcriptomic activity. Given that the relationship between receptor/transporter availability and gene expression is not straightforward (Hansen et al., 2022), the absence of a relationship with either level of observation can be considered complementary evidence. Our exploratory analysis also did not reveal any other molecular function whose transcriptomic activity would suggest a comprehensive relationship between the activation maps and dopamine, norepinephrine, or acetylcholine. This may constrain the strong claim of network-wide neuromodulation, but does not necessarily preclude neuromodulation through other mechanisms, such as more nuanced or dynamic neuromodulatory activity in brain stem or midbrain nuclei. In light of the mixed evidence from pharmacological studies with the ANT (Badgaiyan & Wack, 2011; McCormick, 2022; Reynaud et al., 2019; Thienel et al., 2009), however, it seems reasonable to revise attention network theory regarding the presumed neuromodulation of attention networks.

Finally, our approach is based on the idea that group-level data from three different sources can be combined in a correlational design on the grounds of a cortical parcellation. This methodological approach is widely applied when combing transcriptomic and neuroimaging data (Fornito et al., 2019; Hansen, Markello, et al., 2021; Seidlitz et al., 2020), probing mechanistic hypotheses in clinical neuroscience (Buckner et al., 2008; de Lange et al., 2019; Fornito & Bullmore, 2014; Zhou et al., 2012), and multimodal neuroimaging where the invasive nature of one of assessment methods precludes direct comparisons in the same participants (Heuvel et al., 2015; Scholtens et al., 2014). Despite the limitations of small sample sizes of postmortem brains, constraints in spatial resolution with PET imaging, and unaccounted individual variability, it is important to recognize that the AHBA microarray gene expression maps and group-level receptor/transporter maps still represent state-of-the-art techniques. But even though we used an established null model to reduce spurious influences on our test statistics (Alexander-Bloch et al., 2018; Váša et al., 2018), it needs to be noted that the here presented relationships are correlational. The absence of direct psychopharmacological manipulation or experimental effects on neuromodulators limits conclusions on causality.

### Implications for Attention Network Theory

Since its first conception more than 30 years ago (Posner & Petersen, 1990), attention network theory has established itself as an influential account of how higher cognitive functions such as attention emerges from a network of distributed brain areas (Posner & Dehaene, 1994). By incorporating neuropsychological evidence (Fernandez-Duque & Posner, 2001), modern neuroimaging (Fan et al., 2005; Markett et al., 2014; Petersen & Posner, 2012; Xuan et al., 2016), as well as developmental (Posner et al., 2014), genetic (Fan et al., 2001; Green et al., 2008), and pharmacological data (Marrocco & Davidson, 1998), attention network theory has not only stipulated hundreds of empirical investigations (Arora et al., 2020) but also achieved a level of sophistication that allows for specific hypotheses on the molecular signatures of attention networks. The present findings, however, indicate that some of these predictions do not hold in the proposed way. The ANT activation maps do neither align with the hypothesized distribution of receptor and transporter molecules nor with transcriptomic profiles that would suggest clearly separable networks along molecular lines. Separability and presumed independence of the attention networks is additionally constrained by a high level of spatial dependency between the network maps. Since attention network theory acknowledges interactions between the attention networks (Callejas et al., 2004; Fan et al., 2009; Xuan et al., 2016), it may be reasonable to re-conceptualize the attention networks in terms of their segregation and integration. Future work will also need to re-address the different observational layers and specify how the functional activation maps relate to the underlying brain network (Betzel et al., 2016; Cole et al., 2016; Liu et al., 2022; Markett et al., 2022; Murphy et al., 2020), in order to re-evaluate the presumed independence of attention networks at the neural and neurochemical level, and to specify the presumed neuromodulatory influences on alterting, orienting, and attentional control.

## Acknowledgement

This work was supported by Deutsche Forschungsgemeinschaft, Grant Number: MA-6792/3-1, awarded to Sebastian Markett.

