## SupplementaryMaterials for "Molecular Signatures of Attention Networks"

**Supplementary Materials**

**Appendix A: Genes Products Annotated to Hypothesized Molecular Functions**

| Gene | Full name | Location |
| --- | --- | --- |
| TH | Tyrosine 3-monooxygenase | Chromosome 11, NC_000011.10 (2163929..2171815, complement) |
| DRD2 | D(2) dopamine receptor | Chromosome 11, NC_000011.10 (113409605..113475398, complement) |
| DRD5 | D(1B) dopamine receptor | Chromosome 4, NC_000004.12 (9781634..9784009) |
| DRD4 | D(4) dopamine receptor | Chromosome 11, NC_000011.10 (637269..640706) |
| GPR143 | G-protein coupled receptor 143 | Chromosome X, NC_000023.11 (9725346..9778602, complement) |
| DRD1 | D(1A) dopamine receptor | Chromosome 5, NC_000005.10 (175440036..175444182, complement) |
| SLC6A3 | Sodium-dependent dopamine transporter | Chromosome 5, NC_000005.10 (1392794..1445440, complement) |

*Table A.1.* *Genes annotated to Dopamine Binding (GO:0035240, Organism: Homo sapiens, Gene type: protein coding).*

| Gene | Full name | Location |
| --- | --- | --- |
| ADRB3 | Beta-3 adrenergic receptor | Chromosome 8, NC_000008.11 (37962990..37966599, complement) |
| ADRA2A | Alpha-2A adrenergic receptor | Chromosome 10, NC_000010.11 (111077029..111080907) |
| DRD4 | D(4) dopamine receptor | Chromosome 11, NC_000011.10 (637269..640706) |
| ADRB2 | D(4) dopamine receptor | Chromosome 5, NC_000005.10 (148826611..148828623) |

*Table A.2.* *Genes annotated to Norepinephrine Binding (GO:0051380, Organism: Homo sapiens, Gene type: protein coding).*

| Gene | Full name | Location |
| --- | --- | --- |
| CHRNA3 | Neuronal acetylcholine receptor subunit alpha-3 | Chromosome 15, NC_000015.10 (78593052..78620996, complement) |
| CHRNB4 | Neuronal acetylcholine receptor subunit beta-4 | Chromosome 15, NC_000015.10 (78624111..78661641, complement) |
| CHRNE | Acetylcholine receptor subunit  epsilon | Chromosome 17, NC_000017.11 (4897771..4908677, complement) |
| CHRNA5 | Neuronal acetylcholine receptor subunit alpha-5 | Chromosome 15, NC_000015.10 (78565520..78595269) |
| CHRNA7 | Neuronal acetylcholine receptor subunit alpha-7 | Chromosome 15, NC_000015.10 (32030483..32173018) |
| CHRFAM7A | CHRNA7-FAM7A fusion protein | Chromosome 15, NC_000015.10 (30360566..30393900, complement) |
| CHRNA6 | Neuronal acetylcholine receptor subunit alpha-6 | Chromosome 8, NC_000008.11 (42752620..42768786, complement) |
| CHRNA2 | Neuronal acetylcholine receptor subunit alpha-2 | Chromosome 8, NC_000008.11 (27459756..27479261, complement) |
| CHRNB3 | Neuronal acetylcholine receptor subunit beta-3 | Chromosome 8, NC_000008.11 (42697366..42737407) |
| CHRND | Acetylcholine receptor subunit delta | Chromosome 2, NC_000002.12 (232526160..232536664) |
| CHRNA9 | Neuronal acetylcholine receptor subunit alpha-9 | Chromosome 4, NC_000004.12 (40335333..40355217) |
| CHRNB1 | Acetylcholine receptor subunit beta | Chromosome 17, NC_000017.11 (7445061..7457710) |
| CHRNA4 | Neuronal acetylcholine receptor subunit alpha-4 | Chromosome 20, NC_000020.11 (63343223..63361349, complement) |
| CHRNG | Acetylcholine receptor subunit gamma | Chromosome 2, NC_000002.12 (232539692..232548115) |
| CHRNB2 | Neuronal acetylcholine receptor subunit beta-2 | Chromosome 1, NC_000001.11 (154567778..154580013) |
| CHRNA1 | Acetylcholine receptor subunit alpha | Chromosome 2, NC_000002.12 (174747592..174764472, complement) |
| CHRNA10 | Neuronal acetylcholine receptor subunit alpha-10 | Chromosome 11, NC_000011.10 (3665587..3671384, complement) |

*Table A.3.* *Genes annotated to Acetylcholine-Gated Cation-Selective Channel Activity (GO:0022848, Organism: Homo sapiens, Gene type: protein coding).*

**Appendix B: Detailed Processing Steps**

**Abagen Methods Report**

Regional microarray expression data were obtained from 6 post-mortem brain (1 female, ages 24.0–57.0, 42.50 +/- 13.38) provided by the Allen Human Brain Atlas (AHBA, https://human.brain-map.org; Hawrylycz et al., 2012). Data were processed with the abagen toolbox (version 0.1.3, https://github.com/rmarkello/abagen) using a 219-region surface-based atlas in MNI space.

First, microarray probes were reannotated using data provided by Arnatkeviciute et al. (2019); probes not matched to a valid Entrez ID were discarded. Next, probes were filtered based on their expression intensity relative to background noise (Quackenbush, 2002), such that probes with intensity less than the background in ≥ 50.00 % of samples across donors were discarded, yielding 31,569 probes. When multiple probes indexed the expression of the same gene, we selected and used the probe with the most consistent pattern of regional variation across donors (i.e., differential stability; Hawrylycz et al., 2015), calculated with:

$$\Delta_{s}(p)=\frac{1}{\binom{N}{2}} \sum_{i=1}^{N-1} \sum_{j=i+1}^{N} \rho[Bi(p),Bj(p)]$$

where ρ is Spearman’s rank correlation of the expression of a single probe, p, across regions in two donors *B_i_* and *B_j_,* and N is the total number of donors. Here, regions correspond to the structural designations provided in the ontology from the AHBA.

The MNI coordinates of tissue samples were updated to those generated via non-linear registration using the Advanced Normalization Tools (ANTs, https://github.com/chrisfilo/alleninf). Samples were assigned to brain regions by minimizing the Euclidean distance between the MNI coordinates of each sample and the nearest surface vertex. Samples where the Euclidean distance to the nearest vertex was more than 2 standard deviations above the mean distance for all samples belonging to that donor were excluded. To reduce the potential for misassignment, sample-to-region matching was constrained by hemisphere and gross structural divisions (i.e., cortex, subcortex/brainstem, and cerebellum, such that e.g., a sample in the left cortex could only be assigned to an atlas parcel in the left cortex; Arnatkeviciute et al., 2019). All tissue samples not assigned to a brain region in the provided atlas were discarded. Inter-subject variation was addressed by normalizing tissue sample expression values across genes using a robust sigmoid function (Fulcher et al., 2013):

$$x_{norm}= \frac{1}{1+exp (-\frac{(x-\left\langle x \right\rangle)}{{IQR}_{x}}}$$

where ⟨x⟩ is the median and IQRx is the normalized interquartile range of the expression of a single tissue sample across genes. Normalized expression values were then rescaled to the unit interval:

$$x_{scaled}=\frac{x_{norm}-min(x_{norm})}{max(x_{norm})-min(x_{norm})}$$

Gene expression values were then normalized across tissue samples using an identical procedure. Samples assigned to the same brain region were averaged separately for each donor and then across donors, yielding a regional expression matrix with 219 rows, corresponding to brain regions, and 15,632 columns, corresponding to the retained genes.

**PANTHER Tool Settings and Respective Processing Steps**

- Analysis Type: PANTHER Enrichment Test (Released 20220712)
- Annotation Version and Release Date: GO Ontology database DOI: 10.5281/zenodo.6799722 Released 2022-07-01
- Organism: Homo sapiens
- Annotation Data Set: GO molecular function complete
- Correction: Calculate False Discovery Rate

Using a Mann-Whitney Rank-Sum Test (U-Test), the program calculates a probability to which any genes associated with a particular MF are coincidentally or systematically ranked higher or lower within a gene list. For this purpose, a reference distribution of all genes as well as distributions for each emerging MF is generated, calculating whether the corresponding genes were randomly drawn from the reference distribution (Mi et al., 2013).

**Appendix C. Results of PANTHER Gene Set Enrichment Analyses**

| GO molecular function complete | number | overUnder | pvalue | fdr |
| --- | --- | --- | --- | --- |
| signaling receptor binding (GO:0005102) | 1070 | - | 5.64E-08 | 2.62E-04 |
| structural constituent of ribosome (GO:0003735) | 160 | - | 5.48E-07 | 1.27E-03 |
| active transmembrane transporter activity (GO:0022804) | 313 | + | 5.35E-06 | 8.29E-03 |
| G protein-coupled receptor binding (GO:0001664) | 207 | - | 9.81E-06 | 1.14E-02 |
| cell adhesion molecule binding (GO:0050839) | 487 | - | 1.44E-05 | 1.33E-02 |
| protein serine kinase activity (GO:0106310) | 313 | + | 3.26E-05 | 2.52E-02 |
| ATP binding (GO:0005524) | 1256 | + | 5.36E-05 | 3.56E-02 |
| primary active transmembrane transporter activity (GO:0015399) | 134 | + | 8.38E-05 | 4.87E-02 |

*Table C.1.* *Output of GSEA for the ANT condition alerting.*

| GO molecular function complete | number | overUnder | pvalue | fdr |
| --- | --- | --- | --- | --- |
| structural constituent of ribosome (GO:0003735) | 160 | - | 3.40E-13 | 1.58E-09 |
| DNA binding (GO:0003677) | 1974 | + | 3.18E-11 | 7.40E-08 |
| sequence-specific DNA binding (GO:0043565) | 1313 | + | 3.32E-10 | 5.15E-07 |
| sequence-specific double-stranded DNA binding (GO:1990837) | 1221 | + | 7.95E-10 | 9.24E-07 |
| double-stranded DNA binding (GO:0003690) | 1300 | + | 1.26E-09 | 1.17E-06 |
| transcription cis-regulatory region binding (GO:0000976) | 1177 | + | 2.53E-09 | 1.96E-06 |
| transcription regulatory region nucleic acid binding (GO:0001067) | 1179 | + | 3.00E-09 | 1.99E-06 |
| transcription regulator activity (GO:0140110) | 1541 | + | 3.71E-09 | 2.16E-06 |
| signaling receptor binding (GO:0005102) | 1070 | - | 5.37E-09 | 2.77E-06 |
| nucleic acid binding (GO:0003676) | 3314 | + | 8.69E-09 | 4.04E-06 |
| RNA polymerase II transcription regulatory region sequence-specific DNA binding (GO:0000977) | 1087 | + | 9.96E-09 | 4.21E-06 |
| DNA-binding transcription factor activity (GO:0003700) | 1078 | + | 1.34E-08 | 5.18E-06 |
| DNA-binding transcription factor activity, RNA polymerase II-specific (GO:0000981) | 1038 | + | 1.53E-08 | 5.46E-06 |
| protein serine kinase activity (GO:0106310) | 313 | + | 1.02E-07 | 3.37E-05 |
| cis-regulatory region sequence-specific DNA binding (GO:0000987) | 950 | + | 2.01E-07 | 6.21E-05 |
| G protein-coupled receptor binding (GO:0001664) | 207 | - | 2.24E-07 | 6.50E-05 |
| heterocyclic compound binding (GO:1901363) | 4920 | + | 4.13E-07 | 1.13E-04 |
| ATP binding (GO:0005524) | 1256 | + | 4.67E-07 | 1.21E-04 |
| catalytic activity, acting on a protein (GO:0140096) | 1884 | + | 5.57E-07 | 1.36E-04 |
| organic cyclic compound binding (GO:0097159) | 4971 | + | 6.79E-07 | 1.58E-04 |
| RNA polymerase II cis-regulatory region sequence-specific DNA binding (GO:0000978) | 932 | + | 7.12E-07 | 1.58E-04 |
| protein serine/threonine kinase activity (GO:0004674) | 372 | + | 1.81E-06 | 3.82E-04 |
| structural molecule activity (GO:0005198) | 554 | - | 3.53E-06 | 7.12E-04 |
| adenyl ribonucleotide binding (GO:0032559) | 1310 | + | 4.62E-06 | 8.95E-04 |
| chromatin binding (GO:0003682) | 495 | + | 7.88E-06 | 1.46E-03 |
| oxidoreductase activity (GO:0016491) | 590 | - | 8.86E-06 | 1.58E-03 |
| adenyl nucleotide binding (GO:0030554) | 1322 | + | 9.72E-06 | 1.67E-03 |
| microtubule binding (GO:0008017) | 237 | + | 2.39E-05 | 3.96E-03 |
| phosphotransferase activity, alcohol group as acceptor (GO:0016773) | 588 | + | 2.63E-05 | 4.22E-03 |
| antioxidant activity (GO:0016209) | 66 | - | 3.72E-05 | 5.76E-03 |
| cell adhesion molecule binding (GO:0050839) | 487 | - | 3.81E-05 | 5.70E-03 |
| transmembrane signaling receptor activity (GO:0004888) | 644 | - | 4.54E-05 | 6.59E-03 |
| kinase activity (GO:0016301) | 649 | + | 4.62E-05 | 6.51E-03 |
| receptor ligand activity (GO:0048018) | 275 | - | 4.80E-05 | 6.56E-03 |
| transcription factor binding (GO:0008134) | 545 | + | 5.05E-05 | 6.71E-03 |
| RNA polymerase II-specific DNA-binding transcription factor binding (GO:0061629) | 315 | + | 6.99E-05 | 9.02E-03 |
| cytokine activity (GO:0005125) | 107 | - | 7.08E-05 | 8.89E-03 |
| postsynaptic neurotransmitter receptor activity (GO:0098960) | 55 | - | 8.35E-05 | 1.02E-02 |
| protein kinase activity (GO:0004672) | 492 | + | 8.83E-05 | 1.05E-02 |
| signaling receptor regulator activity (GO:0030545) | 302 | - | 8.85E-05 | 1.03E-02 |
| DNA-binding transcription repressor activity, RNA polymerase II-specific (GO:0001227) | 262 | + | 9.16E-05 | 1.04E-02 |
| signaling receptor activator activity (GO:0030546) | 281 | - | 9.40E-05 | 1.04E-02 |
| molecular transducer activity (GO:0060089) | 793 | - | 9.53E-05 | 1.03E-02 |
| signaling receptor activity (GO:0038023) | 793 | - | 9.53E-05 | 1.01E-02 |
| fatty acid binding (GO:0005504) | 34 | - | 1.02E-04 | 1.05E-02 |
| catalytic activity, acting on a nucleic acid (GO:0140640) | 524 | + | 1.09E-04 | 1.10E-02 |
| DNA-binding transcription repressor activity (GO:0001217) | 266 | + | 1.57E-04 | 1.55E-02 |
| cadherin binding involved in cell-cell adhesion (GO:0098641) | 16 | - | 1.60E-04 | 1.55E-02 |
| copper ion binding (GO:0005507) | 43 | - | 1.77E-04 | 1.68E-02 |
| carboxylic acid binding (GO:0031406) | 121 | - | 1.88E-04 | 1.75E-02 |
| histone binding (GO:0042393) | 228 | + | 1.90E-04 | 1.73E-02 |
| integrin binding (GO:0005178) | 128 | - | 2.05E-04 | 1.84E-02 |
| DNA-binding transcription factor binding (GO:0140297) | 431 | + | 2.24E-04 | 1.97E-02 |
| purine ribonucleoside triphosphate binding (GO:0035639) | 1546 | + | 2.70E-04 | 2.32E-02 |
| glycosaminoglycan binding (GO:0005539) | 158 | - | 3.24E-04 | 2.74E-02 |
| oxidoreductase activity, acting on the CH-NH group of donors (GO:0016645) | 25 | - | 3.72E-04 | 3.09E-02 |
| helicase activity (GO:0004386) | 142 | + | 4.50E-04 | 3.67E-02 |
| metal ion transmembrane transporter activity (GO:0046873) | 339 | + | 4.58E-04 | 3.67E-02 |
| calcium ion binding (GO:0005509) | 530 | - | 4.97E-04 | 3.91E-02 |
| frizzled binding (GO:0005109) | 30 | - | 4.98E-04 | 3.85E-02 |
| transferase activity, transferring phosphorus-containing groups (GO:0016772) | 774 | + | 6.13E-04 | 4.67E-02 |
| peptidase regulator activity (GO:0061134) | 141 | - | 6.18E-04 | 4.63E-02 |
| catalytic activity, acting on RNA (GO:0140098) | 340 | + | 6.37E-04 | 4.70E-02 |
| ATP hydrolysis activity (GO:0016887) | 296 | + | 6.70E-04 | 4.86E-02 |
| bitter taste receptor activity (GO:0033038) | 11 | + | 6.71E-04 | 4.80E-02 |
| ribonucleotide binding (GO:0032553) | 1615 | + | 6.92E-04 | 4.87E-02 |

*Table C.2.* *Output of GSEA for the ANT condition orienting.*

| GO molecular function complete | number | overUnder | pvalue | fdr |
| --- | --- | --- | --- | --- |
| structural constituent of ribosome (GO:0003735) | 160 | - | 5.85E-12 | 2.72E-08 |
| signaling receptor binding (GO:0005102) | 1070 | - | 2.96E-09 | 6.89E-06 |
| structural molecule activity (GO:0005198) | 554 | - | 1.03E-06 | 1.59E-03 |
| transmembrane signaling receptor activity (GO:0004888) | 644 | - | 1.66E-06 | 1.93E-03 |
| catalytic activity, acting on a protein (GO:0140096) | 1884 | + | 2.51E-06 | 2.33E-03 |
| molecular transducer activity (GO:0060089) | 793 | - | 3.67E-06 | 2.84E-03 |
| signaling receptor activity (GO:0038023) | 793 | - | 3.67E-06 | 2.43E-03 |
| G protein-coupled receptor binding (GO:0001664) | 207 | - | 6.08E-06 | 3.53E-03 |
| protein serine kinase activity (GO:0106310) | 313 | + | 2.08E-05 | 1.07E-02 |
| ATP binding (GO:0005524) | 1256 | + | 2.23E-05 | 1.03E-02 |
| cell adhesion molecule binding (GO:0050839) | 487 | - | 2.54E-05 | 1.07E-02 |
| postsynaptic neurotransmitter receptor activity (GO:0098960) | 55 | - | 3.31E-05 | 1.28E-02 |
| integrin binding (GO:0005178) | 128 | - | 5.31E-05 | 1.90E-02 |
| adenyl ribonucleotide binding (GO:0032559) | 1310 | + | 7.34E-05 | 2.44E-02 |
| receptor ligand activity (GO:0048018) | 275 | - | 8.26E-05 | 2.56E-02 |
| microtubule binding (GO:0008017) | 237 | + | 8.64E-05 | 2.51E-02 |
| adenyl nucleotide binding (GO:0030554) | 1322 | + | 1.27E-04 | 3.48E-02 |
| signaling receptor regulator activity (GO:0030545) | 302 | - | 1.50E-04 | 3.88E-02 |
| protein serine/threonine kinase activity (GO:0004674) | 372 | + | 1.50E-04 | 3.68E-02 |
| glutamate receptor activity (GO:0008066) | 26 | - | 1.72E-04 | 3.99E-02 |
| copper ion binding (GO:0005507) | 43 | - | 1.79E-04 | 3.96E-02 |
| cytokine activity (GO:0005125) | 107 | - | 1.81E-04 | 3.83E-02 |
| signaling receptor activator activity (GO:0030546) | 281 | - | 1.91E-04 | 3.86E-02 |
| heterocyclic compound binding (GO:1901363) | 4920 | + | 1.99E-04 | 3.85E-02 |
| neurotransmitter receptor activity (GO:0030594) | 76 | - | 2.48E-04 | 4.61E-02 |

*Table C.3.* *Output of GSEA for the ANT condition control.*

**Appendix D. Description of the MF summarized by GO-Figure!**

**Alerting:**

1. Signaling receptor binding: Signaling receptor binding refers to the interaction between a signaling molecule, such as a hormone, neurotransmitter, or growth factor, and a receptor protein on the surface of a cell. The binding of the signaling molecule to the receptor triggers a series of events within the cell, such as the activation of intracellular signaling pathways or the regulation of gene expression.
2. Structural constituent of ribosome: A structural constituent of ribosome refers to a component that is part of the ribosome's structure. The ribosome is a complex molecular machine found in cells that is responsible for synthesizing proteins. The ribosome consists of several structural constituents, including RNA molecules and proteins, which work together to carry out protein synthesis.
3. Active transmembrane transporter activity: Active transmembrane transporter activity refers to the movement of molecules, such as ions or nutrients, across a cell membrane by a transporter protein. Transporter proteins use energy, typically derived from ATP, to actively transport molecules across the cell membrane. An active transmembrane transporter is one that is currently transporting molecules and is therefore said to have "activity".
4. Protein serine kinase activity: Protein serine kinase activity refers to the ability of a protein kinase to transfer a phosphate group from ATP to a specific amino acid residue, typically serine, on a target protein. Protein kinases play an important role in intracellular signaling by phosphorylating target proteins, which can regulate their activity or localization. The activity of a protein serine kinase refers to its ability to perform this phosphorylation reaction.
5. ATP binding: ATP binding refers to the interaction between adenosine triphosphate (ATP) and a protein molecule. ATP is a high-energy molecule that is used as a source of energy in cells. Many proteins involved in cellular processes, such as enzymes and ion channels, bind ATP as part of their function. The binding of ATP to these proteins can cause structural changes in the protein that activate or regulate its activity. For example, ATP binding to an enzyme can trigger a chemical reaction or ATP binding to an ion channel can open the channel to allow ions to flow into or out of the cell.

**Orienting:**

1. Structural constituent of ribosome: This function refers to a protein's role in being a building block of the ribosome, which is a complex structure within cells that plays a crucial role in protein synthesis.
2. Sequence-specific DNA binding: This function refers to a protein's ability to bind specifically to certain sequences of DNA, which can play a role in regulating gene expression.
3. Double-stranded DNA binding: This function refers to a protein's ability to bind to both strands of DNA at once, which can play a role in DNA replication and repair.
4. Regulatory region nucleic acid binding: This function refers to a protein's ability to bind to specific regions of DNA or RNA that control gene expression.
5. Transcription regulator activity: This function refers to a protein's ability to control the process of transcribing DNA into RNA, which is the first step in making proteins.
6. Signaling receptor binding: This function refers to a protein's ability to bind to receptors on the surface of cells, which can play a role in cellular signaling and communication.
7. Protein serine kinase activity: This function refers to a protein's ability to add a phosphate group to serine residues on other proteins, which can play a role in signaling pathways and regulation of protein activity.
8. ATP binding: This function refers to a protein's ability to bind to the energy-rich molecule ATP, which can play a role in energy metabolism and cellular signaling.
9. Chromatin binding: This function refers to a protein's ability to bind to chromatin, the material that makes up chromosomes and plays a role in regulating gene expression.
10. Antioxidant activity: This function refers to a protein's ability to neutralize harmful molecules called free radicals, which can damage cells and contribute to aging and disease.
11. Transmembrane signaling receptor activity: This function refers to a protein's ability to act as a receptor on the surface of cells, which can play a role in cellular signaling and communication.
12. Fatty acid binding: This function refers to a protein's ability to bind to fatty acids, which are important molecules in energy metabolism and cell membrane structure.
13. Copper ion binding: This function refers to a protein's ability to bind to copper ions, which are important for many enzymes and other proteins in the body.
14. Glycosaminoglycan binding: This function refers to a protein's ability to bind to glycosaminoglycans, which are long chains of sugar molecules that play a role in cell signaling and tissue structure.
15. Oxidoreductase activity, acting on the CH-NH group of donors: This function refers to a protein's ability to transfer electrons from one molecule to another, which can play a role in metabolism and cellular signaling.
16. Helicase activity: This function refers to a protein's ability to unwind and separate strands of DNA or RNA, which is important for DNA replication and other processes.
17. Metal ion transmembrane transporter activity: This function refers to a protein's ability to transport metal ions across the cell membrane, which can play a role in cell metabolism and signaling.
18. Calcium ion binding: This function refers to a protein's ability to bind to calcium ions, which are important for many enzymes and other proteins in the body.
19. Catalytic activity, acting on RNA: This function refers to a protein's ability to catalyze reactions involving RNA, which can play a role in protein synthesis and other processes.

**Control:**

1. Structural constituent of ribosome: This function refers to the role of specific molecules in helping to form the structure of ribosomes, which are responsible for protein synthesis in cells.
2. Signaling receptor binding: This function refers to the ability of certain molecules to bind to specific receptors on the surface of cells, which can initiate a cascade of signaling events that lead to changes in cell behavior or function.
3. Transmembrane signaling receptor activity: This function refers to the ability of certain molecules to act as receptors that span the cell membrane and receive signals from outside the cell, which can then trigger changes in cell behavior or function.
4. Protein serine kinase activity: This function refers to the ability of certain enzymes (called protein kinases) to add a phosphate group to specific amino acids (called serines) on proteins, which can change their activity or localization.
5. ATP binding: This function refers to the ability of certain molecules to bind to the energy-rich molecule ATP (adenosine triphosphate), which can be used to power various cellular processes.
6. Copper ion binding: This function refers to the ability of certain molecules to bind to copper ions, which are important for many enzymes and other proteins involved in various cellular processes, including neurotransmission in the brain.
